# Characterization of the conserved response of angiosperms to hypoxia through transcriptomic meta-analysis

**DOI:** 10.64898/2026.06.14.732220

**Authors:** Sébastien Cabanac, Linh Le Truc Ho, Christophe Dunand, Catherine Mathé

## Abstract

Floods cause significant crop losses worldwide. Plant response mechanisms to flooding have been extensively studied, particularly the ethylene-meditated mechanisms of perception and initiation of the response. However, other mechanisms are often studied more marginally, and it is difficult to determine which are species-specific and which are part of a conserved angiosperm response to hypoxia. Here, we performed a meta-analysis of transcriptomic data under hypoxic or flooding conditions across 11 angiosperm species and identified 259 homologous gene clusters that constitute the core response to hypoxia in angiosperms. These include the previously identified main mechanisms linked to ethylene, as well as numerous novel genes whose role in the hypoxia response is often poorly characterized. In particular, many previously overlooked genes associated with oxidative stress were identified as part of the core response, such as *HRU1, TIP1-2, OZF1*, and *OZF2*. Our results reveal many new candidate genes with strong potential for improving plant resilience to flooding.

## Introduction

Floods are responsible for major losses in agriculture, which relies heavily on the cultivation of angiosperms. Indeed, 17 cultivated angiosperm species provide 90% of human food supply ^1^. During a flood, plants are subjected to waterlogging or, in extreme cases, total submersion. The reduced diffusion of O_2_ and CO_2_ in the water leads to a sudden slowdown in respiration and photosynthesis, hindering energy metabolism. Furthermore, the remaining O_2_ can generate Reactive Oxygen Species (ROS), reactive molecules that can act as a signal but are harmful if they accumulate excessively ^2,3^.

When flooded, plants have mechanisms to perceive and implement a response that allows for ROS scavenging and the establishment of anaerobic metabolism. Some plants also have an escape strategy, elongating until they re-establish contact with the air to allow gas exchange to resume. This energy-intensive process is the opposite of the quiescence strategy, in which a plant limits all growth and unnecessary energy use, favorising post-flooding recovery ^4^.

In all cases, the major mechanism of perception of hypoxia involves the accumulation of ethylene in tissues under hypoxic conditions, which will lead to the stabilization of Class VII Ethylene Response Factor (ERFVII) by a reduction of the Plant Cysteine Oxidase (PCO) N-degron pathway ^5^. Other phytohormones play a more or less limited role in this perception of and response to hypoxia, although their role is less studied ^6^. Similarly, ROS signaling also remains generally less studied. One explanation for this imbalance of information between the role of ethylene and other processes of perception and response to hypoxia could be that, unlike the ethylene-signalling pathway, only a few major genes have been identified for the other hypoxia-related processes. For example, in *A. thaliana, ADH1* and *PDC1* are known to be major regulators of anaerobic metabolism during flooding, while *RBOHD* is important for ROS-mediated signaling ^7^.

In recent years, technical advances have enabled differential transcriptome analysis under flood or hypoxic conditions in many plant species, and a few meta-analyses have combined several results to identify common mechanisms between different plant species, although these are based on a very limited number of angiosperms ^8,9^. Thus, hypoxia response data are available for many angiosperm species, including *Arabidopsis thaliana* ^10–12^, *Brassica napus* ^13^, *Glycine max* ^14^, *Hordeum vulgare* ^15^, *Kalanchoe fedtschenkoi* ^16^, *Lupinus angustifolius* ^17^, *Medicago sativa* ^18^, *Oryza sativa* ^19–21^, *Solanum habrochaites, S. lycopersicum* ^22^ and *Zea mays* ^23^. Data are also available on some aquatic angiosperms, such as *Spirodela polyrhiza* ^24^ or *Kandelia obovata* ^25^, but their response to hypoxia probably differs from terrestrial angiosperms, as suggested by the disappearance of genes involved in ethylene synthesis in some seagrasses ^26^.

We thus took advantage of the large amount of available data to identify mechanisms common to 11 angiosperm species. In addition, we used the different treatment durations between different experiments or within a single experiment to establish the kinetics of the conserved response of angiosperms to hypoxia.

## Results

### Kinetics of the conserved angiosperm response to hypoxia

We retrieved and re-analyzed RNA expression data in response to hypoxia for 14 angiosperm species, including eight dicotyledons (*A. thaliana, B. napus, G. max, K. fedtschenkoi, L. angustifolius, M. sativa, S. habrochaites* and *S. lycopersicum*), and three monocotyledons (*H. vulgare, O. sativa* and *Z. mays*). The proportion of dysregulated genes is quite similar between species, whether for up-regulated genes (mean = 4.91% ± 0.05) or down-regulated genes (mean = 5.37% ± 0.06), with *L. angustifolius* being the strongest exception, although this may be related to differences in experimental protocol. However, the nature of the deregulated genes is rather species-specific, which is marked by a high proportion of orthogroups containing deregulated genes in only a few species (Figure 1). Nevertheless, a small proportion of DEG is shared among a large number of species, and suggests the existence of a “common base” of DEG in response to flooding.

**Figure 1.**
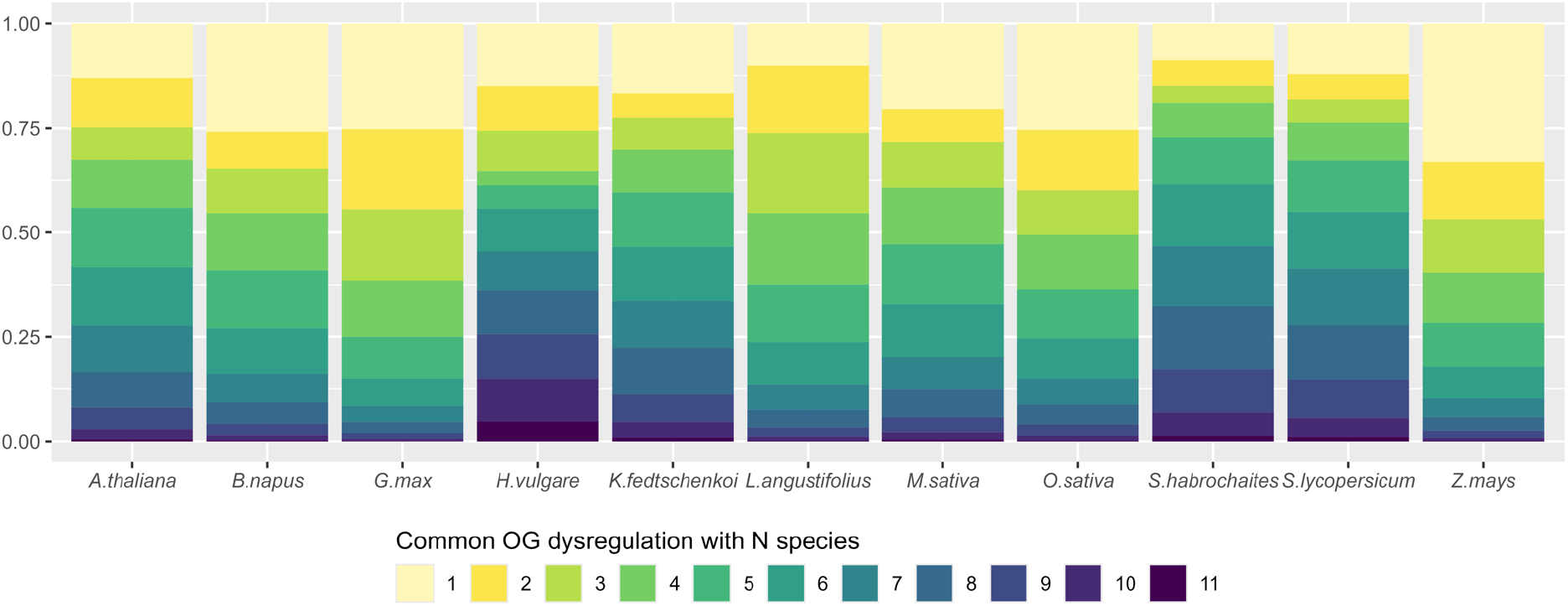
Proportions of orthogroups containing DEGs from a specified number of species.

Since we retrieved data from experiments with different treatment durations for *A. thaliana* and *O. sativa*, and since different kinetics had originally been tested for *Z. mays, G. max* and *B. napus*, we looked at the evolution of the response to hypoxia over time by looking at the proportion of common DEGs between the different time points (Figure 2).

**Figure 2.**
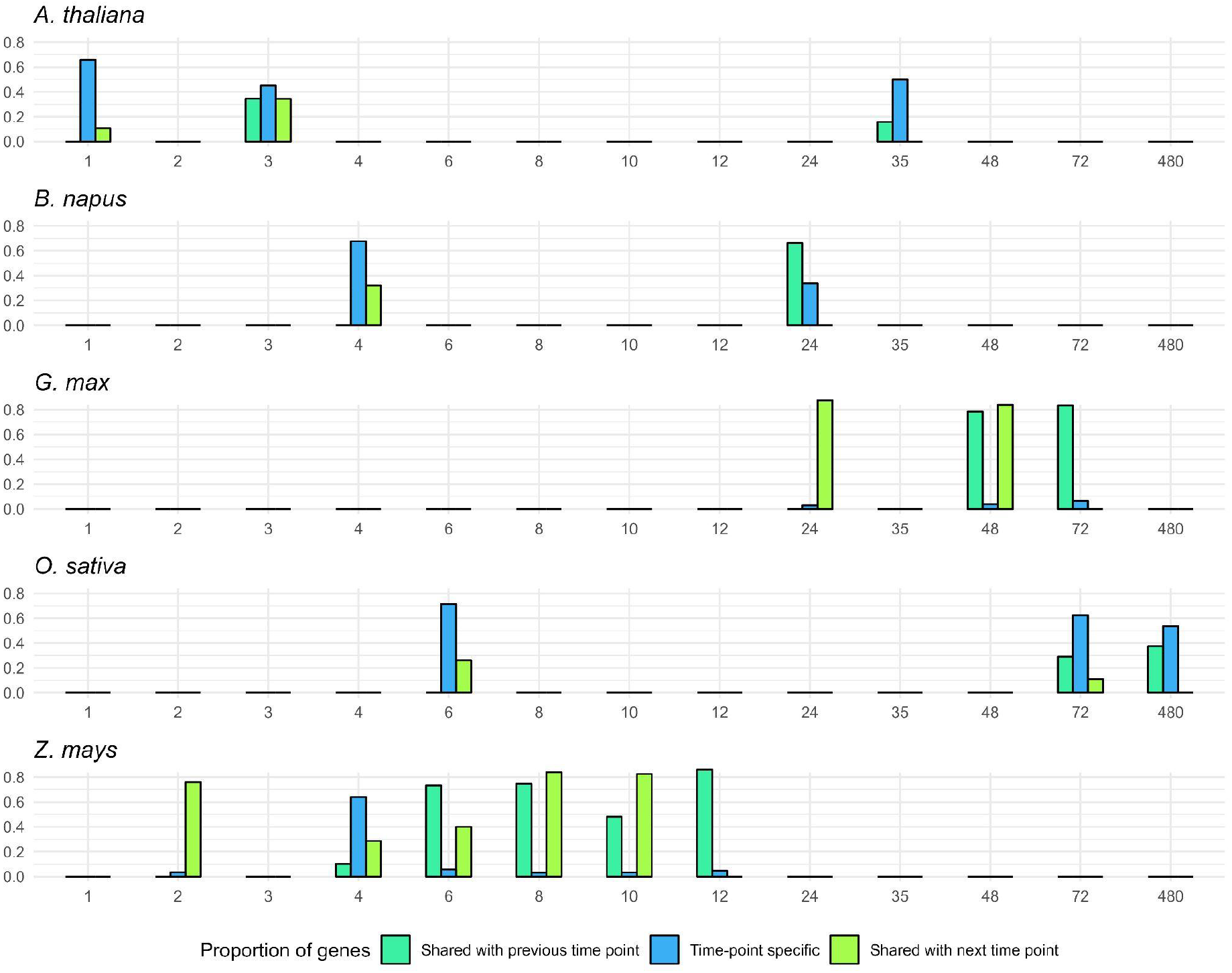
Proportion of DEG shared between different hypoxia durations in *A. thaliana, B. napus, G*.*max, O. sativa* and *Z. mays*.

Data from *A. thaliana* and *Z. mays* show that the DEGs in the first few hours of hypoxia are common across time points, as they share a high proportion of common DEG. However, in both cases, strong specificity is observed at a precise time: 1 hour in *A. thaliana* and 4 hours in *Z. mays*. This shows that the mechanisms of the hypoxia response are activated within the first few hours after the onset of hypoxia and remain active for several hours after. Furthermore, a peak in gene dysregulation is reached within these initial hours, likely corresponding to the perception of hypoxia and activation of the response. Later time points for *B. napus* and *O. sativa* show that the genes dysregulated at longer durations are generally the same as in the first few hours. However, at even later time points, such as 35 hours in *A. thaliana* or 480 hours in *O. sativa*, a higher specificity of DEGs is found, indicating that there may be a response to prolonged hypoxia, but that its kinetics differ between species. Data from *G. max* show that this later response may start as early as 24h, as proportions of shared DEG are very high for every time point past 24 hours.

Most of the results presented so far are based on a small number of time points, and some of the variation between DEF proportion and between time points in *A. thaliana* and *O. sativa* could be explained by different experimental conditions. To address this, we focused on the DEGs common to all the plants studied, further including *H. vulgare* (24h), *K. fedtschenkoi* (8h), *L. angustifolius* (15h), *M. sativa* (336h), *S. habrochaites* (288h), *S. lycopersicum* (288h), and dividing the response to hypoxia into two categories: early (<= 12 h) and late (> 12 h). We focused on orthogroups containing DEGs across all species (common OG), in order to build a set of DEGs that form the conserved angiosperm response to hypoxia (Figure 3).

**Figure 3.**
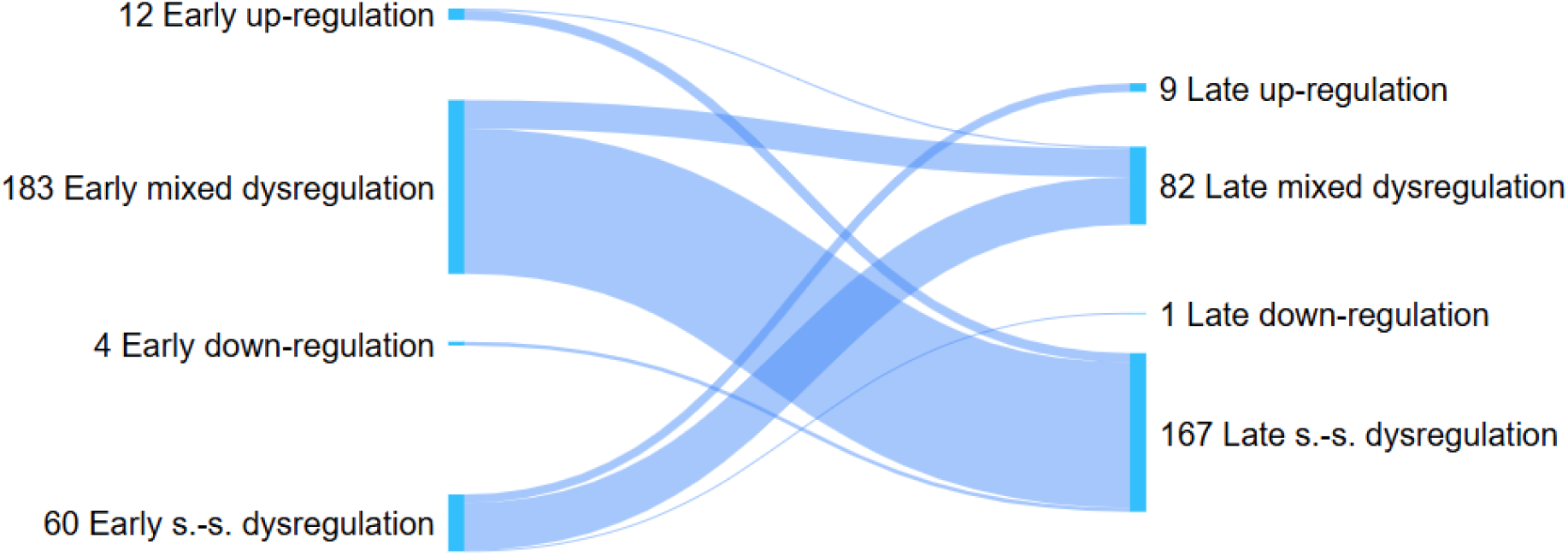
Kinetics of the conserved angiosperm response to hypoxia. Shows orthogroups containing genes that are dysregulated in all species (“common”) and in some species (“s-specific”). “mixed dysregulation” is used to describe orthogroups containing both up-regulated and down-regulated DEGs.

We found 199 common OG at early time points, 91.96% of them having mixed dysregulation. Only 16.08% of these common OGs were found among later-time-points common OGs. Conversely, we found 92 common OGs at later time points, 89.13% of which had mixed dysregulation. 34.78% were also found in early-time-points common OGs. This shows that the conserved response of angiosperms to hypoxia relies on a few homologous genes that are primarily dysregulated in the first few hours of hypoxia. The ability to maintain this dysregulation over longer periods appears to be species-dependent. Conversely, some genes are dysregulated later, and the ability to dysregulate these genes earlier also appears to be species-dependent. In summary, while there is a set of genes commonly found in the conserved response of angiosperms to hypoxia, the kinetics of this response appear to be species-dependent.

### Characterization

Common OGs associated with mixed dysregulation may reveal opposing strategies within plants, or fine-tunings of certain processes. To characterize them, we performed enrichments on *A. thaliana*’s DEGs contained in these common OGs and looked at the most informative results (Table 1).

**Table 1.** Enrichments in Gene Ontology (GO) Biological Process and KEGG pathways obtained with commonly dysregulated orthologous genes from numerous plant species. Only genes from orthogroups with mixtures of up- and down-regulated genes are included. The enrichments were manually filtered to retain only the most informative terms and limit redundancy.

| Category | Identifiers | Description | FDR (Early common OGs) | FDR (Late common OGs) |
| --- | --- | --- | --- | --- |
| GO (Biological Process) | GO:0009733 | Response to auxin | 1E-08 | 5E-04 |
| GO (Biological Process) | GO:0009723 | Response to ethylene | 7E-07 | Not enriched |
| GO (Biological Process) | GO:0009693 | Ethylene biosynthetic process | Not enriched | 9E-06 |
| GO (Biological Process) | GO:0009751 | Response to salicylic acid | 1E-03 | 3E-03 |
| GO (Biological Process) | GO:0009737 | Response to abscissic acid | 1E-03 | Not enriched |
| GO (Biological Process) | GO:0009741 | Response to brassinosteroid | 5E-03 | Not enriched |
| GO (Biological Process) | GO:0010817 | Regulation of hormone levels | 3E-03 | Not enriched |
| GO (Biological Process) | GO:0000302 | Response to ROS | 8E-07 | Not enriched |
| GO (Biological Process) | GO:0006979 | Response to oxidative stress | 5E-06 | Not enriched |
| GO (Biological Process) | GO:0071555 | Cell wall organization | 3E-09 | 3E-02 |
| KEGG pathways | ath04016 | MAPK signaling pathway - plant | 2E-03 | Not enriched |
| KEGG pathways | ath00073 | Cutin, suberine and wax biosynthesis | 8E-03 | Not enriched |
| GO (Biological Process) | GO:0044262 | Cellular carbohydrate metabolic process | 9E-03 | 4E-03 |
| KEGG pathways | ath00010 | Glycolysis / Gluconeogenesis | Not enriched | 3E-04 |
| KEGG pathways | ath00500 | Starch and sucrose metabolism | Not enriched | 3E-02 |
| GO (Biological Process) | GO:0012501 | Programmed cell death | Not enriched | 1E-02 |

Numerous phytohormone-related terms were found among these common OGs. Enrichments in abscisic-acid- and brassinosteroid-related terms were found only in early common OGs, unlike those related to ethylene, auxin, and salicylic acid. This indicates that a fine-tuning of the phytohormone-mediated hypoxia response happens at the onset of a flood, but that this fine-tuning persists only for ethylene, salicylic acid, and auxin during longer flooding. Similarly, our results show that oxidative stress occurs at the beginning of the flood and may be sufficiently reduced to no longer produce enrichment at longer floods. The fact that terms related to oxidative stress are found in these orthogroups with mixed dysregulation may be linked to the dual nature of ROS, which must both be scavenged and serve as a signaling molecule. Finally, terms related to cell wall and carbon nutrition appear in both early and late common OG.

12 common OG contain up-regulated genes from all species at early time points. The *A. thaliana* DEGs contained in these common OGs are primarily involved in phytohormone metabolism, such as *GIM2*, which is involved in ABA catabolism, the negative regulators of the ethylene response *EIN4* and *ETR2*, and the ethylene response transcription factor *ERF1B*. Interestingly, the *HRU1* gene was also found. Its expression is induced by *RAP2-12*, and it was proposed as a link between hypoxia perception and ROS signaling. Another set of genes may be linked to nutrition, and includes the sucrose synthases *SUS1, SUS3, SUS4* and *GDPD1* and *GDPD2*, the former being involved in Pi starvation. Other genes include *STY46, CYP73A5, PSK2, ASK5, ATL6, ATL31, HIR2* and *HIR3*. All of these genes have mixed or species-specific dysregulation over later time points. The same observation is true for common early down-regulated genes, which in *A. thaliana* are *NCED3*, involved in the synthesis of ABA, aquaporin *TIP1-2, BHLH2* and *B3GALT2*.

Conversely, some genes commonly up-regulated at later times are almost all already up-regulated at early times, although this is species-dependent. Conversely, most genes commonly upregulated at later times are almost all already upregulated at early times, although this is species-dependent. These genes are primarily linked to the major mechanism of hypoxia perception and include *PCO1, PCO2, ACO1*, and some ERFVIIs (*HRE2* and *RAP2*.*3*). Interestingly, they also include *HRA1*, a transcription factor that counterbalances the hypoxia response. Other commonly down-regulated genes at later time points include the oxidative-stress-related transcription factors *OZF1* and *OZF2*, the L-lactate dehydrogenase *dl4665w, ATL27, AT1G19530* and *AT5G11090. AVT1J* was the only gene commonly down-regulated at later time points.

## Discussion

Our analyses of a range of angiosperm species have allowed us to determine the conserved response of angiosperms to hypoxia and to characterize its evolution over time. We propose to describe this response as follows: at the onset of a flood, fine-tuning of the hypoxia response is mediated by various phytohormones, including auxin, salicylic acid, abscisic acid, brassinosteroid and ethylene. The ABA response is generally repressed, notably via the upregulation of *GIM2* and the downregulation of *NCED3*, while the ethylene response is negatively regulated, particularly by *EIN4* and *ETR2*, and positively regulated by *ERF1B*. The upregulation of *PCO1, PCO2, ACO1, HRE2, RAP2*.*3*, and *HRA1* also plays a role in fine-tuning the ethylene response, although this process is slower in some species. Up-regulation of these genes is maintained during prolonged flooding, and therefore the ethylene-related response, as well as the salicylic acid and auxin responses, continues during long floods. All of this information is consistent with the scientific literature, including the repression of the ABA response which contributes to survival in flooding, and the major role of ethylene in hypoxia signaling ^6,27^.

The oxidative burst associated with hypoxia is managed in the first hours of flooding, again through fine-tuning of genes linked to ROS homeostasis in order to limit their damage while using them as signaling molecules. To this end, it is possible that the diffusion of ROS, and in particular of H_2_O_2_, is limited by the early down-regulation of *TIP1-2*, one of only two aquaporins involved in the transport of H_2_O_2_ in *A. thaliana. HRU1* also appears to bridge the gap between ethylene-mediated hypoxia signaling and ROS-mediated hypoxia signaling ^28^. Once the initial oxidative burst has passed, the majority of the oxidative stress response subsides. During prolonged flooding, the limited surrounding diffuse oxygen may locally produce oxidative stress, and *OZF1* and *OZF2* could regulate the response to these delayed oxidative stresses, since *OZF1* is involved in the response to oxidative stress ^29^.

Genes related to cell wall remodeling and carbon reserve utilization also appear to be both up- and down-regulated, respectively at the onset of flooding and during prolonged flooding. We cannot definitively conclude that this reflects a distinction between quiescent and escape-strategy plants, as it is difficult to determine whether the varieties used in the original articles employ one or the other of these strategies. It could also reflect fine-tuning of certain processes, such as aerenchyma development or changes in root architecture, this latter point being consistent with the observed hormonal regulations, since it has been noted that ethylene-mediated ABA reduction reactivates adventitious root primordia during flooding ^30^. Nevertheless, across all available data at early time points, *SUS* genes are up-regulated. This is also true later on, although some species exhibit downregulation of certain *SUS*. The major role of the SUS in hypoxia tolerance is known, but the mechanics and their exact role are still much debated ^31–35^. It is possible that this process stops rapidly in quiescent plants, or that it simply does not exist but our data may lack this type of plant to observe it.

Finally, we wish to clarify that this conserved response to hypoxia is certainly not strictly conserved across all angiosperms; but likely reflects the anatomical and environmental characteristics of certain species. For example, the response is probably different in trees, which, due to their height, are less susceptible to total submersion, and the same applies to epiphytic plants. Similarly, this response has probably evolved and differs from what we have observed in aquatic plants, plants adapted to arid environments, or non-photosynthetic parasitic plants. Moreover, although we have focused on angiosperms, part of this response is arguably conserved in other groups that diverged earlier, such as the PCO-related N-degron pathway that exists in *Marchantia polymorpha* and green alga ^36^.

## Conclusion

Analysis of commonly dysregulated orthologs in angiosperms has revealed conserved mechanisms of hypoxia response in these plants. While the major mechanisms of this response have been described previously, notably the ethylene-mediated response, many other genes have been identified. In particular, numerous genes related to the management of oxidative stress and ROS, including *HRU1, TIP1-2, OZF1*, and *OZF2*, have been identified. These genes potentially represent major regulators of the hypoxia response and are good candidates for future studies in plant adaptation to flooding.

## Material and methods

### Data mining

Transcriptomic datasets related to plant hypoxia response were selected using the following query in the NCBI GEO Omnibus database ^37^: (hypoxia) AND “land plants”[porgn: txid3193] NOT “Arabidopsis thaliana”[porgn] NOT “Oryza sativa”[porgn]. Note that *A. thaliana* and *O. sativa* were excluded as we simply retrieved the same experiments as Tamura *et al*. ^9^, although we only kept the experiments carried out on WT. We manually verified each of the results in order to retain only the experiments that actually corresponded to hypoxia assays. Data was downloaded using the fastq-dump 3.1.1 tool. In the end, we used the following datasets (Table 2):

**Table 2.**
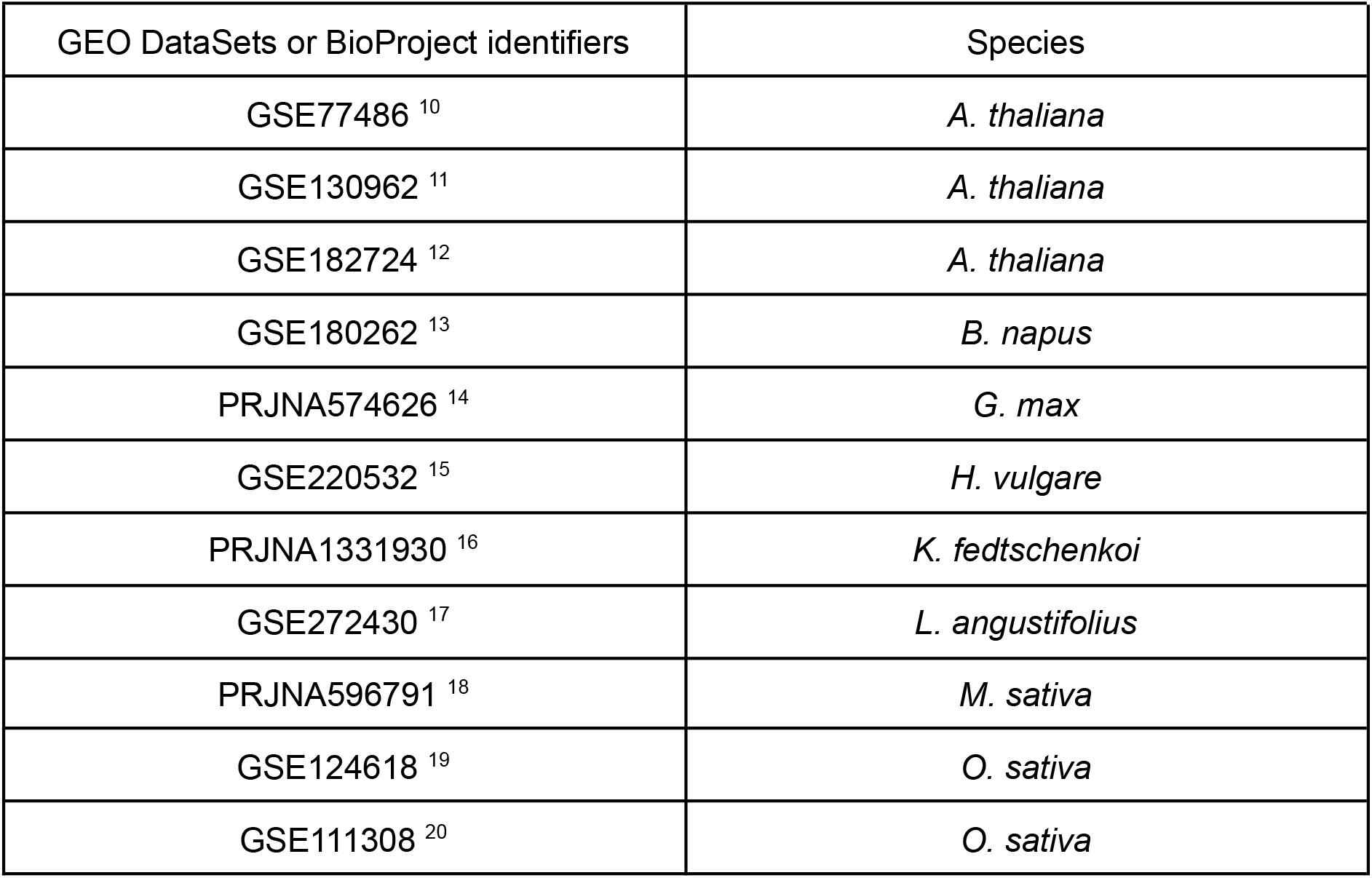
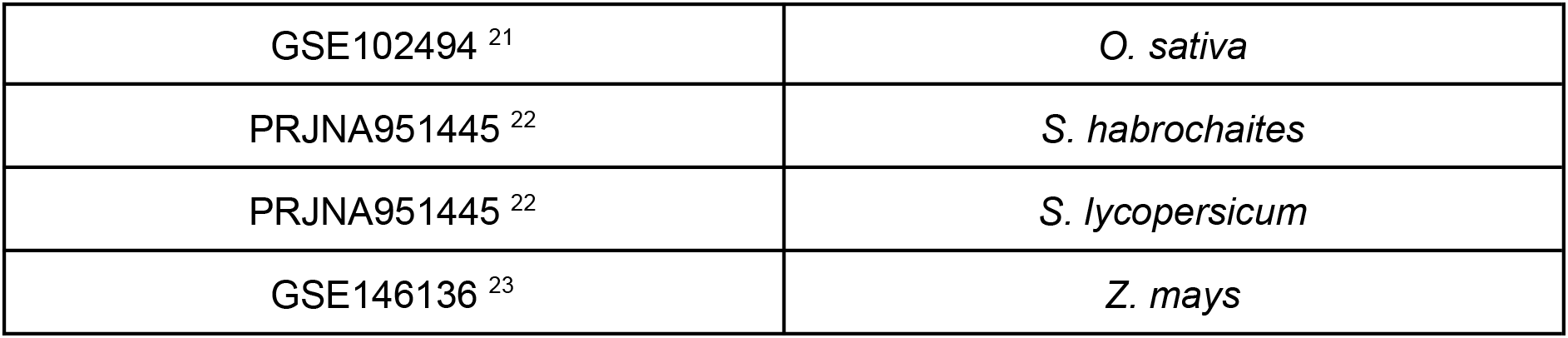
Datasets used in this study.

### DEG selection

Basic quality control was performed with MultiQC v1.35 ^38^, followed by removal of adapters and low-quality bases with TrimGalore v5.0 ^39^. Further quality control was performed on a randomly selected SRR for each species and for each GEO dataset by looking at the mapping rates of the SRR’s reads: 1. on the rRNA SMR “sensitive” v4.3 database using SortMeRNA v4.3.6 ^40^, and 2. on the reference genomes using HISAT2 ^41^ (Table S1) to ensure that any eventual low mapping rates are not due to sample contamination.

RNA quantification was performed by mapping each set of RNA-seq data to reference cDNA FASTA files retrieved from the Ensembl Plants release 62 ^42^ for *A. thaliana, B. napus, G. max, H. vulgare, K. fedtschenkoi, L. angustifolius, O. sativa, S. lycopersicum* and *Z. mays. S. habrochaites* data were downloaded from the Genome Warehouse database ^43^, and M. sativa cDNA file was generated from the Legume Information System’s ^44^ genome and structural annotation using gffread v0.12.7. The completeness of these references was computed beforehand with BUSCO v5.7.1 ^45^ against OrthoDB Embryophyta dataset v10 ^46^ (Table S1).

Alternative transcript quantifications were grouped by gene using the R package tximport v1.30.0 ^47^, and DEG determination was performed with DESeq2 v1.42.1 ^48^, specifying for each experiment the assay SRRs and their corresponding control SRRs (Table S2). When multiple varieties were tested by the original authors who generated the data, we considered a gene as DEG if it appeared as differentially expressed (FDR < 0.05) in the majority of the experiments.

### Orthogrouping

Protein sequence files for all species were downloaded from the same sources as the cDNA sequence files. For each gene, only the longest isoform was retained, and proteins shorter than 80 amino acids were filtered out. Proteins from *Thuja plicata* (v3.1) were downloaded from Phytozome ^49^ and used as an outgroup. Orthogrouping was done using OrthoFinder v2.5.5 ^50^ with -m MSA option.

The type of regulation attributed to each group of homologous genes was defined according to the following rules. An orthogroup was described as up-regulated if it had at least one up-regulated DEG in each of the five species with data on short hypoxic treatments, and at most only one of these species associated with down-regulated genes. For long-term treatments, due to the larger number of species and the lower completeness of the *M. sativa* transcriptome, we described an orthogroup as up-regulated if it contained at least one up-regulated gene in at least seven of the nine species with long treatment data, and at most a single species associated with down-regulated genes. The same approach was used to classify orthogroups as down-regulated. Other orthogroups were considered as having mixed dysregulation.

In addition, orthogroups were classified as commonly dysregulated early if they contained at least one DEG in each of the five species associated with short treatments, and as commonly dysregulated late if they contained at least one DEG in eight of the nine species associated with long treatments. Other orthogroups were considered as having early or late species-specific dysregulation (Table S3).

## Functional enrichment

The functional enrichments were made using the STRING v12.0 web application (https://string-db.org/) ^51^.

## Acknowledgments

We are grateful to the the Genotoul Bioinformatics platform for their support and for providing access to the computing cluster.

